# The Hidden Biology of Wellbeing: A Multi-Omics Analysis Across Genomic, Epigenomic, and Transcriptomic Layers

**DOI:** 10.1101/2025.10.09.681412

**Authors:** Natalia Azcona-Granada, Anne Geijsen, Jenny van Dongen, René Pool, Floris Huider, Pau Badia-i-Mompel, Dirk Pelt, Meike Bartels

**Author notes:** Corresponding author: Mailing address: Department of Biological Psychology, Vrije Universiteit Amsterdam, Amsterdam, Van der Boechorststraat 7, 1081 BT Amsterdam, the Netherlands.

## Abstract

Biological systems are composed of multiple molecular layers, including the genome, epigenome, transcriptome, and metabolome. These layers are interconnected and interact, but are often studied independently. Multi-omics approaches aim to integrate these layers to better understand shared biological processes and their roles in complex traits, such as wellbeing.

This project used data from 2,320 participants from the Netherlands Twin Register (NTR), which includes genome, epigenome, and transcriptome data from the same individuals, providing a rare opportunity to study interactions between omics layers. Multi-Omics Factor Analysis (MOFA) was applied, an unsupervised dimensionality reduction method, to identify latent factors that capture shared variation across omics layers. These factors may reflect underlying biological mechanisms not directly observed in individual omics layers. The phenotypic focus of this study is wellbeing, a multifactorial trait involving emotional, psychological, and social dimensions.

Following MOFA modeling, one latent factor showed to be statistically significantly associated with wellbeing. After studying the latent factor, it mainly seemed to capture epigenetic variation. The association between the latent factor and wellbeing did not remain significant after correcting for relevant covariates. The association was mainly driven by age, a well-known confounder of epigenetic signal.

By adopting a multi-omics framework, this study managed to move beyond traditional single-omics, phenotype-driven approaches and provided a more integrated approach of how biological layers may jointly influence (or not) complex traits like wellbeing.

## Introduction

Biological systems are composed of interconnected molecular layers, traditionally conceptualized through the central dogma: DNA ➜ RNA ➜ protein. However, this traditional model does not reflect the full complexity of how biological systems work since the layers, theoretically, are interconnected and subject to external stimuli (Wolf et al., 2018). This is the reason multi-omics as a field emerged, a discipline in biology that aims to simultaneously characterize and quantify multiple omics layers, including the genome, transcriptome, and metabolome, to provide a more comprehensive view of biological mechanisms (Miller et al., 2023). Each of these omics layers may offer unique insights into a different part of biological processes, helping to understand the bigger picture. The genome provides the blueprint, the epigenome regulates gene expression, the transcriptome captures RNA transcripts, the proteome reflects proteins, and the metabolome comprises small molecules involved in physiological processes.

To get a better understanding of omics for complex traits, many studies have examined associations between individual omics layers and psychological phenotypes (Azcona-Granada et al., 2025; Baselmans, van de Weijer, et al., 2019; van Dongen et al., 2024), integrative data-driven approaches remain rare (e.g., ADHD, aggression; (Hagenbeek et al., 2023; Hubers et al., 2024)). However, since genetic, epigenetic, and transcriptomic layers are biologically linked as parts of cellular regulatory pathways (Fowler et al., 2013), they are expected to share part of their variance. We hypothesize that a factor analysis approach can reveal latent factors that reflect associations among the biological layers, common biological processes, and regulatory mechanisms. These latent factors may, in turn, help explain variation in complex traits such as wellbeing. A latent factor reflects the common variance of one or more omics layers, thereby capturing complex biological interconnections between omics layers. Thus, to obtain a more complete picture, one could investigate the interconnections and overlap among the layers irrespective of a phenotype first, and then look for relations with any phenotype of interest. In this way, the results of the relation between the layers would not be conditioned by the phenotype. In the current study, we take this approach and only in the second step investigate the relation between the omics layers and wellbeing.

Our primary phenotypic focus is on wellbeing, a multifactorial trait involving emotional, psychological, and social dimensions. Wellbeing is defined as the combination of feeling good and functioning well. It includes the experience of positive emotions such as happiness and contentment, as well as the development of one’s potential, having some control over one’s life, having a sense of purpose, and experiencing positive relationships (Huppert, 2009). Like any other complex trait, wellbeing is influenced by multiple biological (and environmental) processes at different levels. The scientific literature has yielded a plethora of studies investigating the multifactorial determinants of wellbeing across these different levels of analysis. These levels include the genome, epigenome, and transcriptome, which will be discussed in depth here, given their relevance for this study.

For example, twin studies consistently show that genetic factors account for about 30–50% of the variance in wellbeing, with the remaining variance largely due to unique environmental influences (Bartels, 2015; Røysamb et al., 2018, 2023). Molecular genetic studies have shown that wellbeing is a polygenic trait. The first genome-wide association study (GWAS) identified 13 genome-wide significant loci (Okbay et al., 2016), while a Multi-Trait Analysis of GWAS (MTAG) increased the number of loci to 49 (Turley et al., 2018). A more recent multivariate GWAS identified 304 independent genome-wide SNPs associated with the wellbeing spectrum, which includes life satisfaction, positive affect, neuroticism, and depressive symptoms (Baselmans, Jansen, et al., 2019).

Although no other biological layer has been studied in such detail as the genome, interesting findings from other layers exist. An EWAS study found two CpGs significant for wellbeing after Bonferroni correction and six applying FDR instead (Baselmans et al., 2015). A MWAS analysis, which detects genetically driven CpGs associated with wellbeing by combining GWAS and mQTL summary statistics, found 913 CpG methylation–trait associations with wellbeing (Baselmans, Jansen, et al., 2019). More recently, an EWAS meta-analysis based on blood DNA methylation found no significant CpGs for wellbeing (Bartels et al., 2025).

Transcriptomics, serving as the intermediary stage between DNA and protein expression, remains a relatively underexplored area in relation to wellbeing. Baselmans et al. (2019) found 97 transcript–trait associations with wellbeing in their GWAS. In addition, transcriptomic differences have been observed for phenotypes related to wellbeing, such as major depressive disorder (MDD) (Sforzini et al., 2024; Sokolov et al., 2024), and other phenotypes related to psychopathology (e.g., schizophrenia) (Wagh et al., 2021).

To date, no study has applied a multi-omics approach to wellbeing. Multi-omics approaches have been applied to other behavioral phenotypes. For depression, for instance, integrating data across omics layers has revealed complex biological mechanisms—including immune, metabolic, and gut-brain interactions—and helped identify molecular signatures linked to different subtypes and onset patterns (Grant et al., 2022; Habets et al., 2023; Hernández-Cacho et al., 2025; Stolfi et al., 2024).

Summarizing, wellbeing is a complex, multifactorial trait likely influenced by the interaction between various biological layers. To disentangle these relationships, we apply Multi-Omics Factor Analysis (MOFA) (Argelaguet et al., 2018) to genomics, epigenomics, and transcriptomics data. This method extends the principles of traditional factor analysis to the multi-omics context. Just as regular factor analysis identifies latent factors that capture shared variation across observed variables, MOFA extracts low-dimensional factors that summarize variation both within and across omics layers. These factors are interpretable through loadings of the features, meaning that for each feature, it is quantified how much it contributes to the factor. The factors can reveal underlying biological processes or shared sources of variability across layers. By integrating multiple omics layers, rather than analyzing each in isolation, we aim to determine whether this joint modeling approach improves the prediction and understanding of the biological mechanism explaining differences in wellbeing. In addition to evaluating predictive performance, we also interpret the meaning of the factors by investigating the respective feature loadings to gain insights into the molecular architecture of wellbeing and how different biological systems interact to shape it.

## Methods

This is a pre-registered project (https://osf.io/rpjmn).

### Sample

Participants included in this study are part of the Netherlands Twin Register (NTR), a population-based national register of twins and their families with extensive data collection on, amongst other things, mental health, personality, lifestyle, and demographics (Ligthart et al., 2019). Participants have been recruited since 1986, and questionnaires have been sent out every 2–3 years. In addition to questionnaires, the biological samples used in this project were collected between 2004 and 2008. The questionnaires, biological sampling, processing, and storage are described in previously published work (Ligthart et al., 2019; Willemsen et al., 2010).

The data used in this study were obtained from the NTR after meeting the requirements determined by the NTR Data Sharing Committee. All procedures performed in studies involving human participants were in accordance with the ethical standards of the institutional and/or national research committee and with the 1964 Helsinki Declaration. Data collection was approved by the Central Ethics Committee on Research Involving Human Subjects of the University Medical Centers Amsterdam. Informed consent was obtained from all participants in the study.

A sub-sample from the NTR biobank (*n* = 2,320) was selected with complete data for all three omics layers (genome, epigenome, and transcriptome) assessed in this project. For the analysis of the overlap between the omics layers and wellbeing, 1888 individuals had overlapping omics, covariate, and wellbeing data. The mean age was 33 (*SD* 12.8, range 17–79 years). The ratio of men/women was 632 (33.5%) / 1256 (66.5%). In this study, we utilized data from adult participants of surveys 6 (2006–2007), 8 (2009–2010), 10 (2013–2014), and 14 (2019–2020). In cases where multiple wellbeing measures were available, we selected the one closest in time to the biological sample collection.

### Measures

#### Biological layers

In 2004, the Netherlands Twin Register (NTR) started a large-scale biological sample collection in twin families to create a resource for genetic studies on health, lifestyle, and personality. Between January 2004 and July 2008, adult participants from NTR research projects were invited into the study. During a home visit between 7:00 and 10:00 am, fasting blood and morning urine samples were collected. Fertile women were bled on day 2–4 of the menstrual cycle, or in their pill-free week. Biological samples were collected for DNA isolation, gene expression studies, creation of cell lines, and biomarker assessment. At the time of blood sampling, additional phenotypic information concerning health, medication use, body composition, and smoking was collected. The collection of DNA and RNA has been described in previous publications (Ligthart et al., 2019; Willemsen et al., 2010). All biological data were scaled to a Z-score before further integration to make the different biological layers comparable.

##### Genetic data

Quality control of the genotype data is explained in detail in the Supplementary Material. Genotype data included single-nucleotide polymorphisms (SNPs) assessed using the Affymetrix 6.0, Illumina General Screening Array, and Affymetrix AXIOM-NL arrays. Genotype quality control included the removal of SNPs with poor imputation quality, low minor allele frequency (0.01), or those not complying with the Hardy-Weinberg equilibrium (0.0001). SNPs were aggregated into/annotated to genes using the ENSEMBL GRCh38 (hg38) reference panel containing gene and enhancer data (Smedley et al., 2009), accessed via BioMart. We used the *map_snp_to_gene* function from the R package *snpsettest* (Joo & Himes, 2023) to map SNPs to genes based on their physical position within the genome. This function assigns SNPs to genes within a specified window (1kb upstream and 0.5kb downstream), enabling downstream gene-level association analyses. Based on the annotated data, SNP values were aggregated into a single gene score value by averaging over all SNPs belonging to that gene. Since the SNP notation (0,1,2) represents the number of alternative alleles – not necessarily the effect allele – from the BioMart reference panel, a higher score means a higher number of alternative alleles for that gene (González Silos et al., 2022). To normalize the data, a logarithmic (log_e_) transformation was applied to the overall gene scores for 30,817 genes mapped from 1,086,331 SNPs, not including X, Y, or mitochondrial genes.

##### Methylation data

The epigenomic data were assessed with the Infinium HumanMethylation450 BeadChip Kit. Genomic DNA (500 ng) from whole blood was bisulfite-treated using the ZymoResearch EZ DNA Methylation kit (Zymo Research Corp, Irvine, CA, USA), following the standard protocol for Illumina 450k microarrays, as performed by the Department of Molecular Epidemiology at the Leiden University Medical Center (LUMC), The Netherlands. Subsequent steps (that is, sample hybridization, staining, and scanning) were performed by the Erasmus Medical Center microarray facility, Rotterdam, The Netherlands. QC and normalization of the methylation data were carried out with pipelines developed by the Biobank-based Integrative Omics Study (BIOS) consortium, as was previously described (Sinke et al., 2019) and has been described in detail previously (van Dongen et al., 2016). In short, samples were removed if they failed to pass all five quality criteria of MethylAid (van Iterson et al., 2014), if samples had incorrect relationships (omicsPrint; (van Iterson et al., 2018)), or sex mismatches (DNAmArray; (Min et al., 2018)). Methylation probes were set to missing in a sample if they had an intensity value of zero, a bead count <3, or a detection p-value>0.01. DNA methylation probes were excluded if they overlapped with a SNP or Insertion/Deletion (INDEL), mapped to multiple locations in the genome, or had a success rate <0.95 across all samples. Missing methylation β-values were imputed (missMDA; (Sinke et al., 2019)), and only one sample was included for NTR participants, with longitudinal DNA methylation data were excluded. Based on the standard deviation of each CpG, a filter for the top 50% most variable CpGs was applied, resulting in a subset of 205,577 CpGs. Different thresholds for the variance of methylation data have been used before, mainly to reduce the computational burden. We selected the top 50% most variable CpGs based on the standard deviations in our sample (Hagenbeek et al., 2025).

##### Transcriptome data

The generation of the gene expression arrays and data in the NTR was described before (Jansen et al., 2014; Wright et al., 2014). Briefly, peripheral venous blood samples were drawn in the morning (7:00—11:00 a.m.) after an overnight fast. Within 20 min of sampling, heparinized whole blood was transferred into PAXgene Blood RNA tubes (Qiagen) and stored at −20°C. The frozen PAXgene tubes were shipped to Rutgers University and the DNA Repository (RUCDR, http://www.rucdr.org). RNA was extracted by the Qiagen Universal liquid handling system, as per the manufacturer’s protocol. RNA quality and quantity were assessed by Caliper AMS90 with HT DNA5K/RNA LabChips. Samples were hybridized to Affymetrix U219 array plates (GeneTitan), which contain 530,467 probes for 49,293 transcripts. Array hybridization, washing, staining, and scanning were carried out in an Affymetrix GeneTitan System per the manufacturer’s protocol. Twin pairs were randomized over the sample plates. Expression data were required to pass standard Affymetrix Expression Console quality metrics before further undergoing quality control. Samples were excluded if they showed sex inconsistency. Log2 transformation and quantile normalization were applied to the gene expression data. The process resulted in 44,241 transcripts, including probes for X and Y chromosomes.

#### Wellbeing

Wellbeing was assessed as a continuous trait combining three measures. Quality of Life (QoL) was assessed with the Cantril ladder, which invites participants to indicate the step of the ladder at which they place their lives in general on a 10-point scale (10 being the best possible life, and 1 the worst possible life) (Cantril, 1965). Satisfaction with life was assessed with the Satisfaction With Life Scale (SWLS, (Diener et al., 1985)). The SWLS consists of five items answered on a seven-point scale ranging from 1=“strongly disagree” to 7=“strongly agree.” An example item is: “I am satisfied with my life.”. Subjective happiness was measured with the Subjective Happiness Scale (SHS, (Lyubomirsky & Lepper, 1999)). The SHS consists of four items, which are rated on a seven-point scale ranging from 1=“strongly disagree” to 7=“strongly agree.” An example item is “On the whole, I am a happy person”.

Satisfaction With Life, Subjective Happiness, and Quality of Life were summarized into a single score for wellbeing using a latent factor score, as was done previously (Azcona-Granada et al., 2025; Bartels et al., 2013). The sum scores for QoL, SWLS, and SHS were first transformed into Z-scores to avoid scaling issues when creating the latent factor score. In each wave, a single factor score for wellbeing was built on all available wellbeing measures using structural equation modeling (Bartels et al., 2013). We used full information maximum likelihood when constructing the factor score, which can handle missing values on indicator variables. However, to limit the number of missing values per participant, only those with at least two out of three wellbeing measurements per wave were included. For more details on the construction and reliability of the latent factor score, see an earlier publication (Azcona-Granada et al., 2025).

#### Covariate data

The covariates included in this study were age at blood sampling, sex, Body Mass Index (BMI) at blood sampling, smoking status at blood sampling, time between collection of the survey data and the collection of the biological data, white blood cell count, D, batch effects, and 20 genetic ancestry principal components. Smoking status was coded as ex-smoker, never smoker, or current smoker. White blood cell count includes % neutrophils, % lymphocytes, % monocytes, % eosinophils, % basophils, and the total white blood cell count from both the epigenome and transcriptome samples. D is a transcriptome array batch measure (Wright et al., 2014). In the epigenetic data, the batch effect was assessed with the sample plate (dummy-coded) and the array row number.

### Statistical analyses

For the proposed analyses, we applied an unsupervised multi-factor analysis method called Multi-Omics Factor Analysis (MOFA) (Argelaguet et al., 2018), which was performed in R (version 4.5) with the MOFA2 package (Argelaguet et al., 2018). MOFA is a powerful computational method for integrating and analyzing multi-omics data, enabling the identification of latent factors that explain variability across different biological layers. In this study, our input data for the fitting of the MOFA model consisted of gene variants, CpGs, and transcripts (Figure 1A, left). In the first step, we used these data to explore different numbers of latent factors to uncover meaningful biological patterns among the omics layers. After identifying these factors, individuals’ factor scores were saved. In step two, to test whether wellbeing or any of the other covariates was associated with our inferred factors, we computed Pearson correlations. We correlated the saved factor scores with wellbeing and with all the covariates for each omics layer, such as age, sex, BMI, smoking status, time difference, white blood cell count, D, batch effects, and genetic ancestry principal components using Pearson correlations.

**Figure 1.**
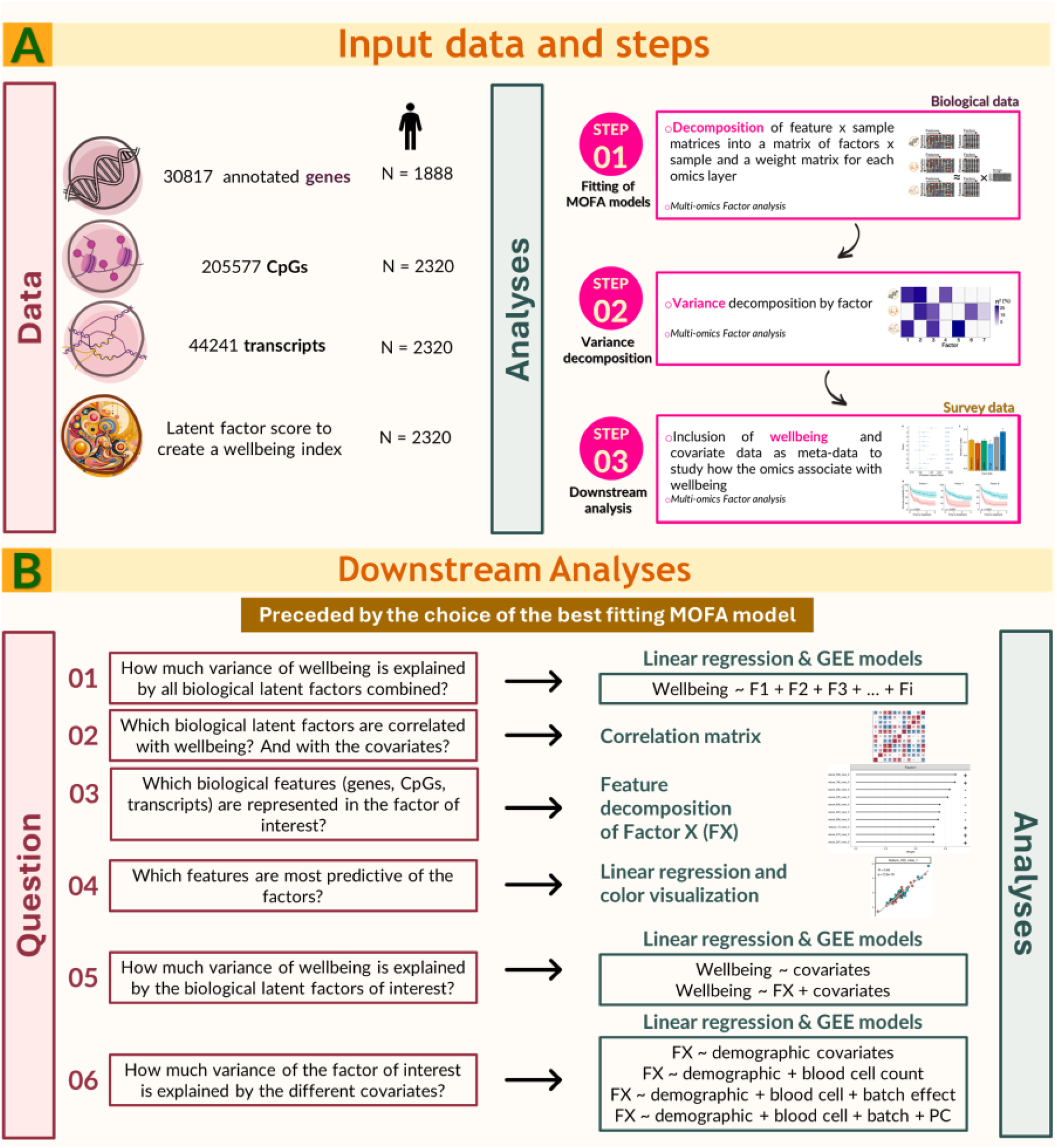
A) Overview of MOFA input data (left) and the steps of the analyses (right). B) Downstream analyses. On the left the question aimed to answer, on the right the analyses performed for that question. F denotates factor; FX denotates the significant factors. From step 03 onwards, only the significant factors are kept.

For the fitting of the MOFA model – the first step – we used the largest available sample (2,320 individuals). In the second step, also referred to as downstream analyses, we included the meta-data (i.e., wellbeing and covariate data) for all 2,320 individuals, regardless of whether they had some missing data. For example, only 1,888 individuals had data for wellbeing, but all 2,320 individuals were included in the downstream analyses because MOFA can handle missing data.

#### Fitting of the MOFA model

MOFA was fitted with the R package MOFA2 (*MOFA2*, n.d.) using default settings and model options, with layers scaled to account for different measurement ranges. The model was optimized using 10 and 20 factors, and fitting was executed in a high-performance computing environment.

#### Downstream analyses

After fitting the model, the variance explained by the 10-factor model and the 20-factor model was assessed to evaluate which model significantly captures more biological information. The variance explained for each factor by each biological layer (i.e., genome, epigenome, transcriptome), as well as the total sum of all the variance explained in all the factors by each biological layer, was extracted to understand the contribution of each biological layer to the model. Factor scores per individual for the model that captured the most biological information were extracted and saved for further downstream analyses.

For a general understanding of the explained variance in wellbeing by all the biological layers, we regressed wellbeing on all the factors together and report the *R*^2^ value (Figure 1B; question 01).

Next, the variables for the downstream analyses, i.e., wellbeing and the covariates, were correlated with the factor scores with Pearson correlations to test whether any of the identified factors was significantly correlated with wellbeing (Figure 1B; question 02). Correlations between the continuous variables (e.g., latent factors resulting from the model, wellbeing, age, time difference, BMI, white blood cell count, and D) were calculated in *lavaan* (Rosseel, 2012). Standardized estimates from regressions with Generalized Estimated Equation (GEE) models with the R package *gee* (Carey, 2023) were calculated as a proxy for the correlation when one or both of the variables were categorical (sex, smoking status). Both in *lavaan* and the GEEs, we corrected for the clustering of families. A Benjamini-Hochberg correction (also called False Discovery Rate (FDR)) was used as the multiple testing correction method (Rouam, 2013). We aimed to identify the factors correlated with wellbeing and to explore the relationships these factors have with other relevant variables. Importantly, only the factor(s) that showed a significant bivariate (unadjusted) correlation with wellbeing were investigated in all the following steps.

Once we established which (if any) of the latent factors were associated with wellbeing, downstream analyses of the feature composition and loadings of the biological layers on those factors were performed with the same MOFA package in R (MOFA2). A brief description of all the downstream analyses is shown in Figure 1B. For the downstream analyses of the significant factor(s), we performed a more in-depth analysis of the feature contribution of all three omics layers to that factor (Figure 1B; question 03). Factor loadings of the optimal factor model were plotted following the pipeline of MOFA downstream analysis (*MOFA+: Downstream Analysis in R*, n.d.) and colored based on the level of the wellbeing scores and any covariate that was also significantly correlated with the factor of interest (Figure 1B; question 04). This visualization allowed us to explore whether samples grouped by the significant variables/covariates, e.g., wellbeing, age, etc., formed distinct clusters when plotting factor scores against feature values, suggesting a relationship between the latent factor and the phenotypic variable.

Next, to identify if the factor(s) of interest were independently correlated with wellbeing or if the association was driven by covariates, hierarchical GEE linear regressions were conducted with wellbeing as the outcome (Figure 1B; question 05). The first model included only the covariates; in the second model, each factor was added. We compared the R^2^ of the model including only the covariates with the R^2^ of a model including the factor(s) of interest (statistically significantly associated with wellbeing), and calculated the incremental R^2^, representing the incremental effect of the factor on wellbeing beyond the covariates.

Finally, to understand the extent to which the variance captured by the factor(s) of interest overlapped with the variance in the covariates, GEE models were calculated regressing the factor scores on all the significant covariates (Figure 1B; question 06). We compared the R^2^ of four models: A) including only the demographic covariates; B) including the demographic covariates and the blood cell count covariates; C) model B + the batch effect covariates; and D) model C + the genetic PCs. The regression model used to calculate all the incremental R^2^ were general linear models since the variance explained is not affected by family relatedness, and therefore, there is no need to correct for family structure.

## Results

### Fit of MOFA models

After fitting both the 10-factor MOFA model and the 20-factor MOFA model, we proceeded to compare them. To understand the structure of each fitted MOFA model, we compared them by investigating correlations between the factor scores of the 10-factor and 20-factor models. The first five factors of both models were strongly correlated between both models (r = 0.85 – 1). Factors 6 to 10 of the 10-factor model were strongly correlated with factors 7, 8, 11, 12, and 14, respectively, from the 20-factor model (for more details, see Supplementary Figure 1). To understand which one of the models explained the most variance and, therefore, is most appropriate for downstream analyses, we extracted the variance explained by each fitted model. The variance explained by each model in the biological layers is presented in Table 1. For the variance explained per factor per biological layer, see Supplementary Tables 1 and 2.

**Table 1.**
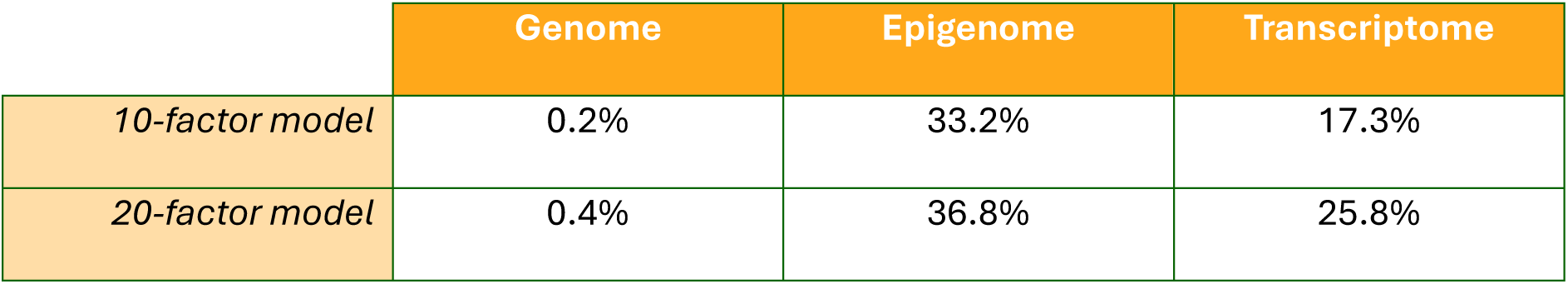
Relative variance of each omics layer – Genome, Epigenome, and Transcriptome – explained by the two different models – 10 and 20 latent factors – trained by MOFA. The total variance explained per omics layer in MOFA does not sum to 100% because MOFA typically explains only a subset of the total variance (i.e., the shared structure). The remainder is residual (unexplained) variance, which could be due to noise, omics-specific structure, or irrelevant variation.

|  | Genome | Epigenome | Transcriptome |
| --- | --- | --- | --- |
| 10-factor model | 0.2% | 33.2% | 17.3% |
| 20-factor model | 0.4% | 36.8% | 25.8% |

The 10-factor model explained 50.7% of the variance observed in the biological data. The 20-factor model explained 63.0% of the variance. The remaining 49.3% and 37.0%, respectively, are residual (unexplained) variances, which could be due to noise, omics-specific structure, or irrelevant variation (systematic variation unrelated to the primary biological question, such as technical artifacts or confounding effects). Since the 20-factor model showed a marked increase in variance explained – especially for the transcriptome and (in a relative sense) the genome, this was the chosen model to continue with the downstream analyses. Supplementary Table 2 shows that the last factors explain basically no variance. Absolute correlations between the 20 factors were very small, ranging from 0 to 0.14.

### Downstream analyses

First, for a general understanding of the biological variance explained, we calculated the variance of wellbeing explained by all the latent factors in the 20-factor MOFA model. The totality of all the biological layers captured by the latent factors in the MOFA model was R^2^ = 0.02 (R^2^ adj = 0.01). Then, we investigated which of the specific factors were contributing to wellbeing.

The variables included were wellbeing and the covariates age, sex, BMI, smoking status, time difference, % neutrophils, % lymphocytes, % monocytes, % eosinophils, % basophils, total white blood cell count, array row, sample plate, D, and 20 genetic ancestry principal components.

After correlating all the latent factors of the model with the meta-data variables, factor 16 significantly correlated with wellbeing after FDR correction (*r* = 0.09, p_adj = 0.0027) (Figure 2A).

**Figure 2.**
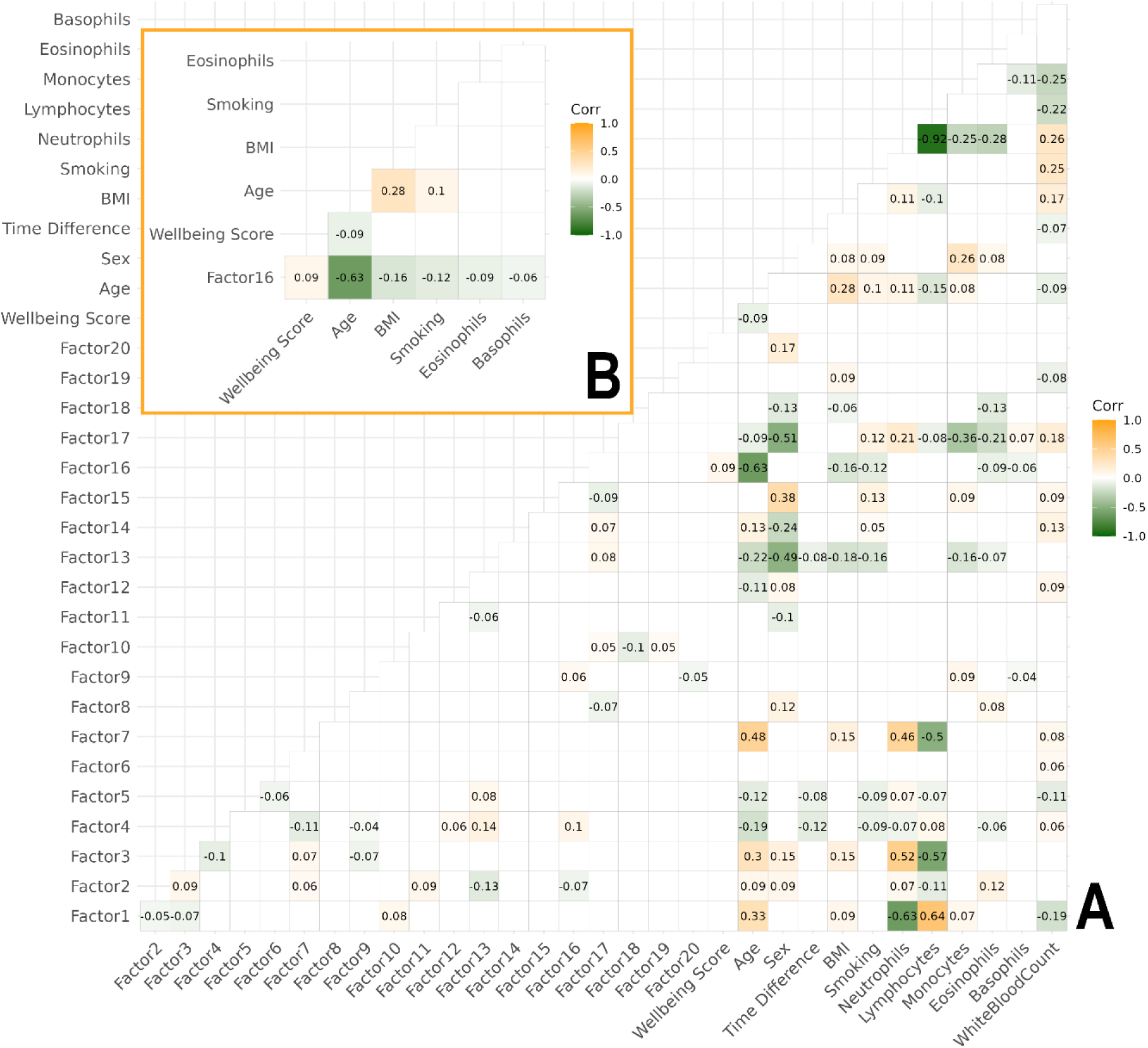
A) Correlation matrix between the latent factors from the MOFA model and the meta data included in the model. Only shown are the significant correlations after FDR correction. B) Correlation matrix between Factor 16, wellbeing, and significant covariates for Factor 16. Note: only the significant correlations after FDR correction are shown. Technical covariates (genetic PCs, epigenetic, and transcriptomic batch effects) are not shown in the figure.

Besides wellbeing, Factor 16 was also significantly correlated with the covariates age, BMI, smoking status, % eosinophils, and % basophils. For ease of interpretation, Figure 2B zooms in on Factor 16 with the statistically significantly associated variables.

To understand which features and biological layers are most strongly represented by Factor 16, we performed a feature decomposition of this factor. Ranking the features by their contribution i.e., loading strength, we found that the top 16,271 features originated from the epigenome layer (i.e., CpG sites). Since this factor is mostly explained by epigenome data, we focus on CpGs in the following results. For illustrative and readability purposes, the first 20 CpGs, as well as their respective feature loadings in Factor 16, are presented in Figure 3. To see the genes associated with these CpGs, see Supplementary Table 3.

**Figure 3.**
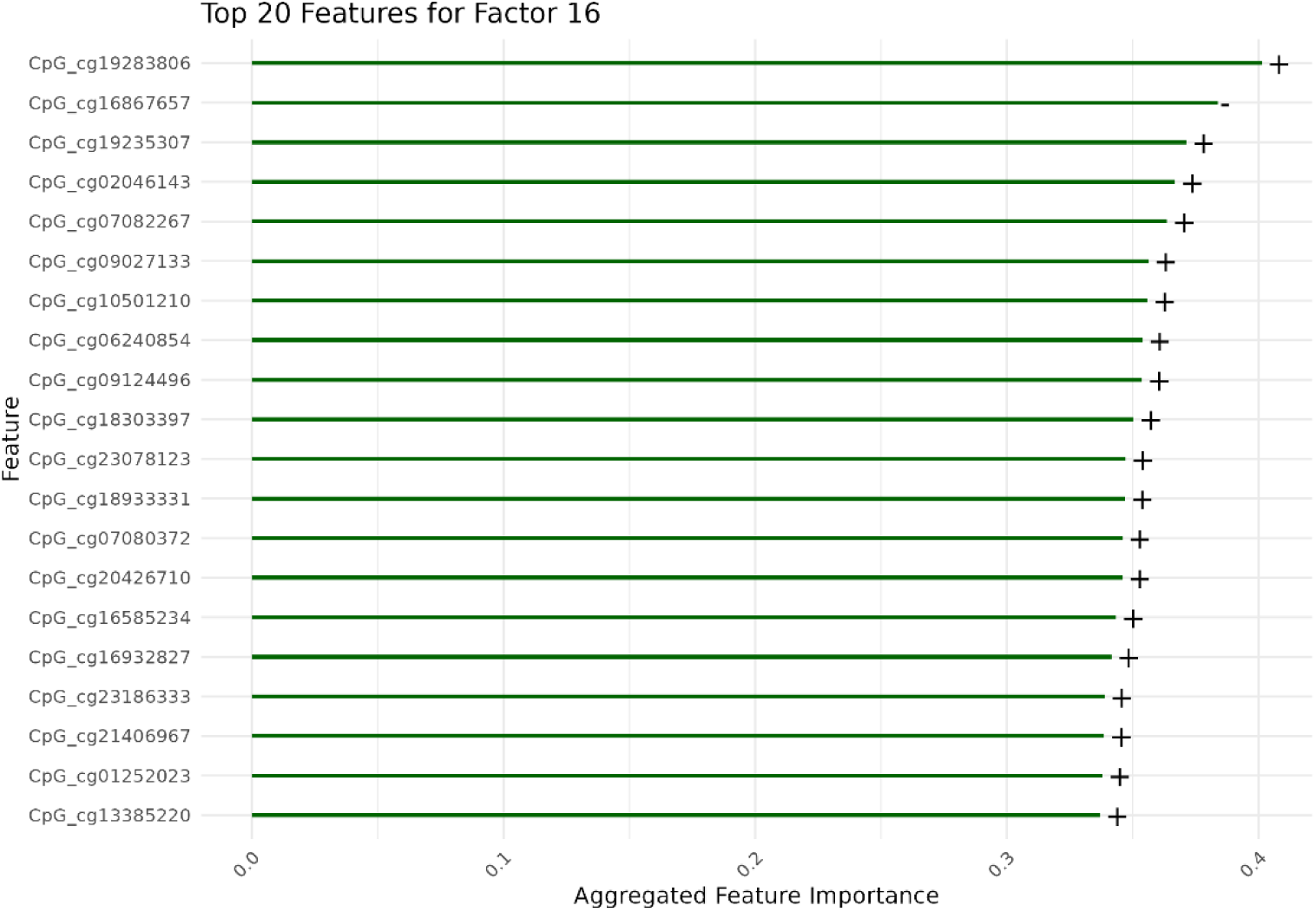
First 20 features with the largest loading for Factor 16 and the direction of their loading (+: positively correlated with factor 16; –: negatively correlated with factor 16).

As part of the MOFA downstream analyses, we evaluated to what extent the selected CpGs are associated with the latent factor scores by assessing the proportion of variance in the factor explained by these features (*R*²). This allows us to quantify the relevance of the CpGs to the factor and assess their predictive contribution. For illustrative purposes, we only focus on the three most predictive CpGs. These first three CpGs had an *R*^2^ ranging from 0.55 to 0.59 (*P* < 2.2e-16). In Figure 4, we show a scatter plot of CpGs associated with Factor 16, where each point represents an individual, with the color indicating individuals’ scores on wellbeing or one of the significant covariates (age, BMI, smoking status, % eosinophils, and % basophils). This visualization allows us to assess whether any of these covariates show distinct clustering patterns in relation to the factor. Notably, visible clustering emerged only for age, suggesting a stronger relationship between age and the factor compared to the other variables included, as previously seen in the correlation matrix.

**Figure 4A.**
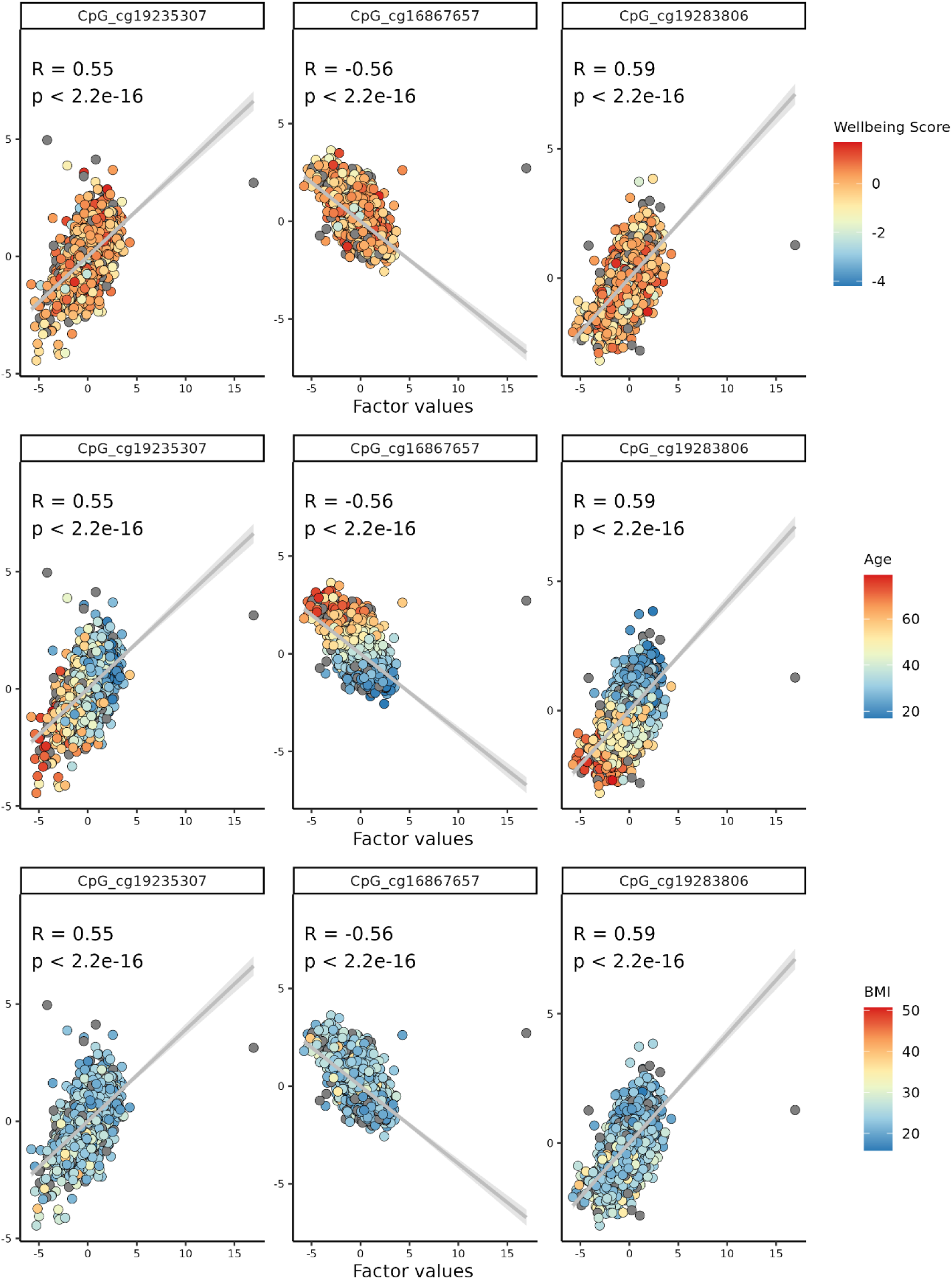
Factor prediction from the first three CpGs colored by wellbeing score (top), age (middle), and BMI (bottom).

**Figure 4B.**
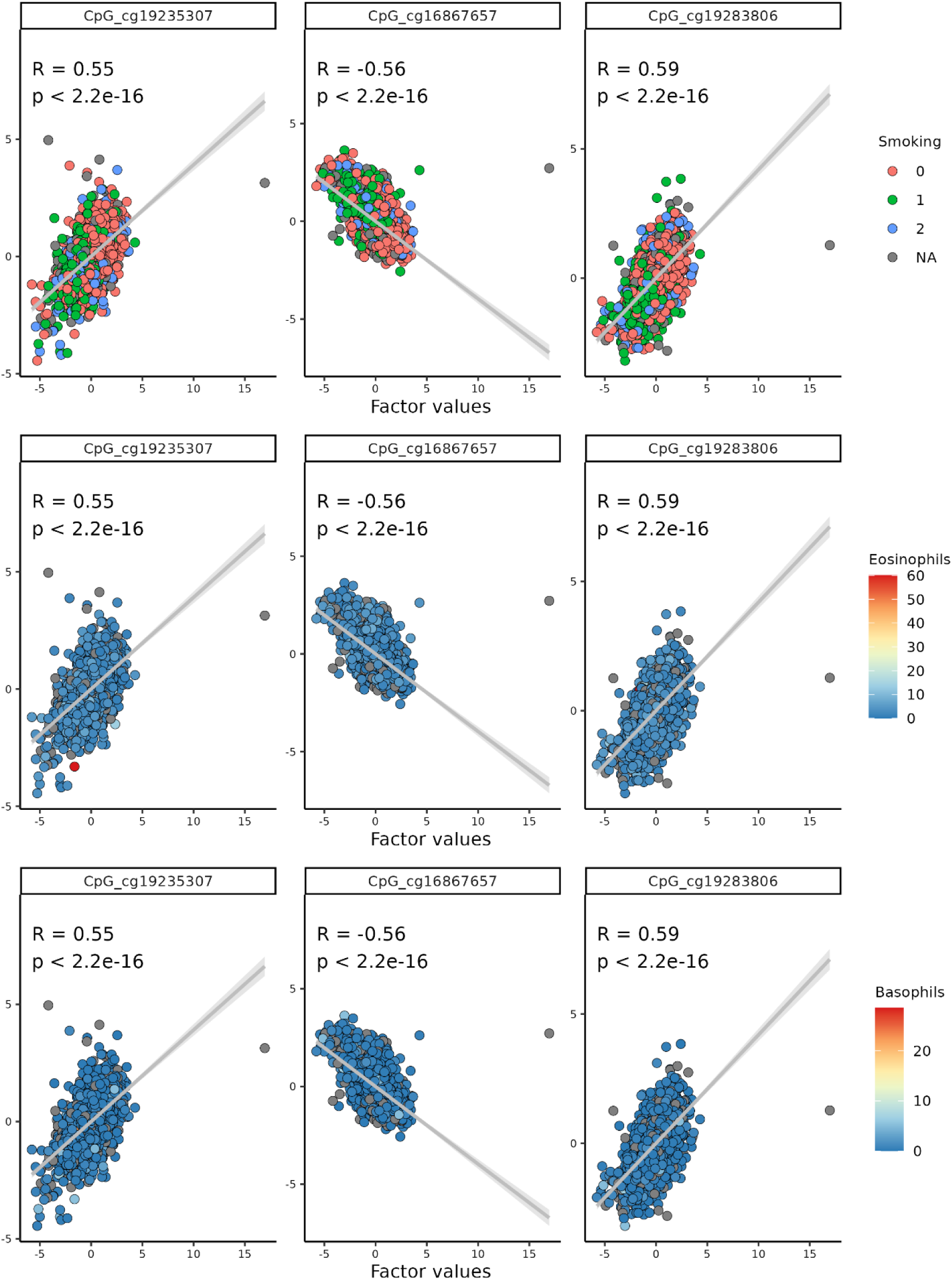
Factor prediction from the first three CpGs colored by smoking (top), % eosinophils (middle), and % basophils (bottom).

Since no visual clustering of wellbeing was present, we conducted hierarchical GEE linear regressions to determine whether Factor 16 was independently associated with wellbeing, or whether the association was driven by the covariates (see Figure 1B, question 05). In Model 1, wellbeing was predicted by the covariates alone. In Model 2, both the covariates and the factor of interest were included as predictors. In Model 1, the *R*^2^ was 0.0601 (R^2^ adj = 0.03), meaning that 6.01% of the variance in wellbeing was explained by the covariates age, BMI, smoking status, % eosinophils, % basophils, array sample plate, array row, and 20 genetic PCs. In model 2, the R^2^ was 0.061 (R^2^ adj = 0.03), meaning that the incremental R^2^ (ΔR^2^) added by Factor 16 was ΔR^2^ = 8.57 e-04 (ΔR^2^ adj = 0.0003). In this Model 2, there was no significant effect of Factor 16 on wellbeing (Factor 16 on Model 2: β = 0.04, *P* = 0.20).

To get insights into which variables were driving values on Factor 16, we additionally compared the incremental *R*^2^ of four models with Factor 16 scores as the outcome (see Figure 1B, question 06): A) Factor 16 predicted by only the demographic covariates, i.e., age, BMI and smoking status; B) adding the percentage of white blood cell count, namely % eosinophils and % basophils; C) adding the batch effects, namely array row number and sample plate; and D) adding the genetic principal components (20 PCs). The variance explained for group A was calculated to be *R*^2^ = 0.38 (*R*^2^ adj = 0.38), adding group B the Δ*R*^2^ = 0.00 (ΔR^2^ adj = 0.00), for group C was ΔR^2^ = 0.07 (ΔR^2^ adj = 0.05) and for group D was ΔR^2^ = 0.00 (ΔR^2^ adj = 0.00).

## Discussion

In behavioral sciences, factor analysis and related multivariate techniques have been widely used to study behavioral traits, especially in the context of personality and psychopathology (Awogbemi et al., 2025; Watts et al., 2023). However, in our study we use Multi-Omics Factor Analyses (MOFA) for a behavioral phenotype such as wellbeing. This study aimed to understand whether biological layers could be linked through shared latent factors and whether this overlap is linked to wellbeing. Indicated by strong loadings of a single layer on each factor and small correlations between the factors, no clear overlap between the biological omics layers was found. Only one of the 20 latent factors (Factor 16) capturing the biological variance was significantly correlated with wellbeing after FDR correction. However, no significant associations between this latent factor and wellbeing remained after correction for the covariates. The epigenome features, i.e., CpGs, explained most variance, both overall and in the aforementioned factor associated with wellbeing. Despite the absence of significant associations between biological latent factors and wellbeing, these analyses offer a novel perspective on the biological underpinnings of wellbeing and highlight the complexity of integrating multi-omics data in complex trait prediction. Given the lack of similar studies, direct comparison of our results to previous work is not possible yet, and replication is needed to confirm our findings.

While other omics layers did not show strong associations with wellbeing in our analyses, the genetic contribution to wellbeing has been well-documented. As previously mentioned, when the genome is studied as a single layer, multiple results have been published (Bartels et al., 2022; van de Weijer et al., 2022). However, as mentioned before, in the current study, the genetic layer explained very little variance in the biological data. This difference with the single omics studies may be caused by little variation between individuals in the genetic data, or maybe if any variance is present, it may be washed out by the variance in the other omics layers. It is also possible that if the genetic effect sizes are smaller than the rest, the sample size required is larger to observe them. Supporting the notion that genetic effects on wellbeing can be subtle, a recent study by de Vries et al. (2024) performed a GWAS by subtraction to isolate “pure” wellbeing, identifying only one significant SNP per construct and observing a notable decrease in SNP heritability (de Vries et al., 2024). Together, these findings suggest that while genetics contributes to wellbeing, the signal may be weak or context-dependent, especially when “pure” wellbeing is studied.

In the present study, the contribution to the variance explained by the transcriptome was larger than the variance explained by the genome. However, the transcriptome showed weak associations compared to the epigenome. This is partly in line with previous literature. The transcriptome explains a significant portion of variance in biological studies (Bhattacharya et al., 2021; García-Nieto et al., 2022). These studies make the transcriptome analysis a central component of multi-omics strategies for understanding complex biological systems and disease mechanisms. However, based on this literature, we expected larger effects of the transcriptome layer, which could have been masked by the number of features contained in the epigenome layer.

As previously mentioned, the epigenome was the largest contributor to Factor 16. The isolated association between the epigenome and wellbeing has been previously investigated (Bartels et al., 2025). Bartels et al. (2025) found that an aggregated methylation score – calculated as the sum of individual CpG effects associated with wellbeing –significantly predicted wellbeing levels in the NTR sample. The same study found significant CpGs for wellbeing in the meta-analysis of 13 cohorts (Bartels et al., 2025). Of note, in our prior EWAS meta-analyses, we adjusted the covariates that emerged in our current analysis as being associated with factor 16, as is common practice in EWAS analyses.

A review of multi-omics studies integrating genetic, transcriptomic, and epigenetic data also shows that epigenetic changes are often associated with phenotypic outcomes – such as gray matter atrophy in depression, aging, and longevity (Mavromatis et al., 2023; Zheng et al., 2024). These studies suggest that epigenetic modifications can show strong associations with biological traits, sometimes independently of, or in addition to, genetic variation. It is also well-known that the epigenome is the best predictor of ageing, smoking and cellular composition (Klopack et al., 2022). In this context, our finding that the strongest signal appears to come from the epigenome is not unusual.

As seen in previous studies, when using biological data, it is essential to correct for covariates. We found the previously established association between Factor 16 and wellbeing to vanish after controlling for age, BMI, smoking status, % eosinophils, % basophils, batch effects, and genetic ancestry PCs. Since there was a clear relation between the factor and age, as seen in the correlations and visually shown in Figure 5, it appears that any relation between the factor and wellbeing was actually driven by age. This result is not surprising, given how closely related the epigenome and age are (Kabacik et al., 2022; Mareckova et al., 2023). Epigenomic changes, among which are DNA methylation, are connected with other hallmarks of aging (Booth & Brunet, 2016; Wang et al., 2022) such as genomic instability, cellular senescence, and altered intercellular communication. The epigenomic changes are key in the aging process, but aging is also key in the epigenetic changes, meaning the association is bidirectional. This means that when the epigenome is studied in relation to outcomes such as wellbeing, age is an important covariate to take into consideration. While research on epigenetic clocks is currently booming (Teschendorff & Horvath, 2025), our study shows it requires careful consideration of age as a confounder.

In this study, age is not only associated with Factor 16, but also significantly correlated with wellbeing (r = –0.09). This negative association is in line with previous published analyses (Baselmans et al., 2018). Results from Hansen and Blekesaune et al. (2022) indicated negative changes in advanced age across wellbeing measures. They found loss of health, a partner, and friends as robust predictors of declining wellbeing (Hansen & Blekesaune, 2022). It shows that wellbeing changes over the lifetime and therefore it should be considered.

It is worth mentioning the different strengths and limitations we faced during this project. MOFA is a complex analytical approach that detects subtle patterns across latent dimensions. This is the first study applying MOFA to wellbeing with a relatively large sample size (N = 2,320). In our effort to keep the input data as raw as possible to retain the full scope of information, we may have ended up with an unbalanced data structure, particularly due to the high number of CpGs compared to genes or transcripts. However, further reducing the epigenomic layer would have resulted in a significant loss of information. As for the other layers – genomic and transcriptomic – additional data were simply not available to a sufficient extent (see below). Despite these limitations, the amount of data we were able to include and integrate into the model is noteworthy.

Data from other omics layers are available for the NTR, and studies have been published on their association with wellbeing individually. For example, on the metabolome level, even though no significant metabolite remained after multiple testing correction, the results suggested a role of lipid metabolism in wellbeing (Azcona-Granada et al., 2025). Despite acknowledging the importance of metabolites for wellbeing, the number of features (143 metabolites) was too small relative to the other three layers for inclusion in the current study.

Future studies would benefit from larger datasets with increased participant numbers to enhance statistical power. Additionally, achieving a more balanced representation of features across each omics layer would improve model robustness. Although proteomic data are not currently available in the NTR biobank, integrating additional biological layers may reveal overlapping signals across omics, as seen before for other phenotypes, like depression (Habets et al., 2023). One possible reason why genome-wide association signals were less apparent in the present study is that SNPs were aggregated at the gene level; examining SNPs directly, where computationally feasible, may uncover effects otherwise obscured by aggregation. Moreover, since all omics layers except genomics are inherently tissue dependent, relying on blood-derived measures may limit the detection of associations that could emerge in other tissues. With the modeling pipeline now established and having demonstrated promising results, future research in the NTR could explore extending this framework to other phenotypes, particularly those with distinct biological signatures such as depression, stress, or aggressive behavior.

In summary, although we hypothesized that a factor analysis approach may reveal latent factors that reflect associations among the biological layers with wellbeing, no clear shared biological pattern was observed across the included biological layers. The latent factor most strongly associated with wellbeing was primarily driven by epigenomic variation. However, this association was no longer significant after adjusting for covariates, and the relation seemed to be mostly driven by age. To increase understanding of the biological underpinnings of complex traits like wellbeing, future research will require the integration of larger and more balanced datasets, both in terms of omics layers and sample size.

## Acknowledgments

We warmly thank all participating twins in the Netherlands Twin Register who dedicated part of their time to make research possible as well as everyone involved in the collection of the data and data management. This study is funded by an NWO-VICI grant (VI.C.211.054, PI Bartels). Data collection for this study has been funded by NWO large investment grant (NTR: 480-15-001/674), ZonMW Addiction program (31160008), ERC Consolidator Grant (WELL-BEING; grant 771057), NWO 575-25-006, Genotype/phenotype database for behavior genetic and genetic epidemiological studies (ZonMw Middelgroot 911-09-032); and ERC Advanced, 230374.

