## Supplementary Material for "The Hidden Biology of Wellbeing: A Multi-Omics Analysis Across Genomic, Epigenomic, and Transcriptomic Layers"

**Appendix**

**Supplementary Genetic QC information.**

The participants were genotyped as part of the larger NTR study, with QC, cleaning and imputation performed at the cohort level before extraction of the subset with DNA methylation data.

We used version 18 of the NTR genotyping data. Genotyping was conducted on three SNP array platforms: Affymetrix 6.0 (N=10,377), Affymetrix Axiom (N=3,536), and Illumina GSA NTR array (N=20,060). Genotype calling followed the manufacturer protocols and white papers. DNA samples were checked for gender mismatches, heterozygosity and Identity by Descent (IBD) mismatches in comparison to the known family structure. Each sample needed to have a call rate of at least 90% and at least 80% of the genotypes needed to be present on each chromosome. SNP quality control was based on the following filters applied in each platform: call rate over 95%, Hardy-Weinberg equilibrium (HWE) p-value over 0.0001, minor allele frequency over 0.01 and Mendelian as well as genotyping error rate less than 1%. This led to the final set of 3,644 individuals with 534,405 SNPs on Axiom, 9,049 individuals with 537,992 SNPs on Affymetrix 6.0 and 16,276 individuals with 481,898 SNPs on Illumina GSA, a total of 28,969 individuals. After strand and name alignment of the data, removing palindromic SNPS with a MAF>0.30, the genotype data were imputed per platform with Beagle 5.4 against the 1000 Genomes Phase 3 set and the HRC 1.1 reference panel (McCarthy et al., 2016). After imputation, the imputed data were merged into a single dataset using bcftools. The HRC imputed data were used for the current GoDMC analyses. The 1000Genomes imputed data were only used to predict ancestry and remove ancestry outliers, as described in the next paragraph, prior to running the GoDMC pipeline.

Twenty 1000 Genomes projected principal components (PC) for the genotype data were calculated based on SNPs that passed quality control (QC) and were present in one of the three platforms from the 1000 Genomes imputed data (as the overlap between platforms is too small to take only genotyped SNPs). SNPs were then filtered to have MAF>0.05, HWE p > 0.001, call rate > 0.98, Mendelian error rate < 1% and imputation info>=90%. These SNPs were subsequently pruned for linkage disequilibrium (LD) with Plink version 1.9 (option –indep 50 5 2) and SNPs in long range LD blocks were removed as described in (Abdellaoui et al., 2013). This left 110,558 SNPs for PC analysis. From the 1000 Genomes reference panel all samples with the same SNPs were selected and then merged with the NTR data. Subsequently PCs were calculated in the 1000 Genomes set and then projected upon the NTR data with the smartpca software to identify and remove ancestry outliers.

**Supplementary Figure 1. Correlation matrix between the models**

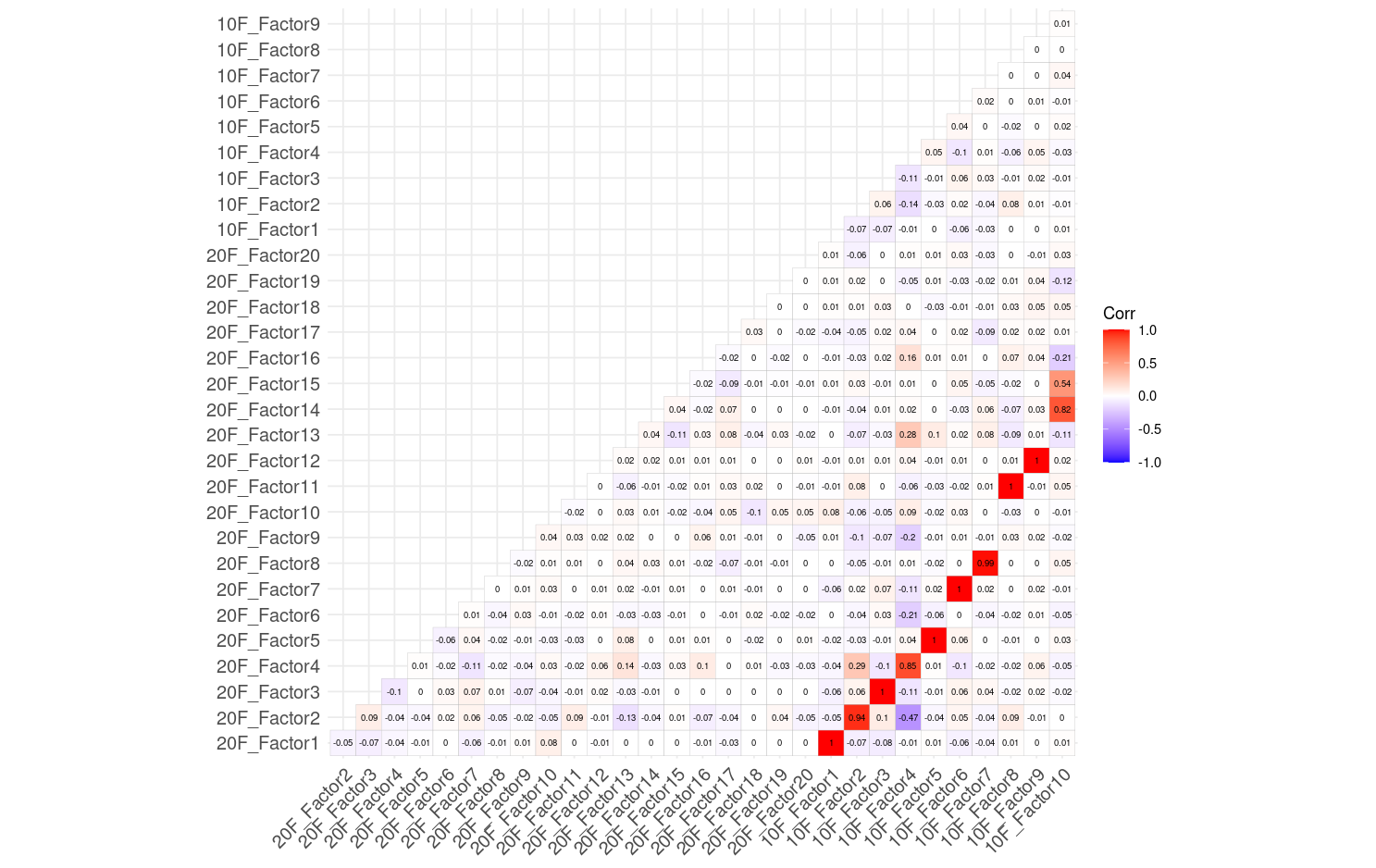

**Supplementary Table 1. Variance explained by omics layer per factor in the 10 factors model.**

|  | Genome | Epigenome | Transcriptome |
| --- | --- | --- | --- |
| Factor 1 | 0.02 | 12.24 | 0.58 |
| Factor 2 | 0.01 | 0.01 | 9.46 |
| Factor 3 | 0.02 | 7.35 | 0.65 |
| Factor 4 | 0.02 | 0.09 | 6.48 |
| Factor 5 | 0.01 | 4.52 | 0.05 |
| Factor 6 | 0.02 | 3.15 | 0.41 |
| Factor 7 | 0.02 | 2.56 | 0.03 |
| Factor 8 | 0.01 | 1.30 | 0.02 |
| Factor 9 | 0.02 | 1.14 | 0.02 |
| Factor 10 | 0.01 | 0.79 | 0.11 |

**Supplementary Table 2. Variance explained by omics layer per factor in the 20 factors model.**

|  | Genome | Epigenome | Transcriptome |
| --- | --- | --- | --- |
| Factor 1 | 0.02 | 12.26 | 0.57 |
| Factor 2 | 0.01 | 0.00 | 10.34 |
| Factor 3 | 0.025 | 7.37 | 0.60 |
| Factor 4 | 0.02 | 0.03 | 5.11 |
| Factor 5 | 0.01 | 4.50 | 0.04 |
| Factor 6 | 0.00 | 0.00 | 3.72 |
| Factor 7 | 0.02 | 3.15 | 0.40 |
| Factor 8 | 0.02 | 2.50 | 0.03 |
| Factor 9 | 0.00 | 0.01 | 2.29 |
| Factor 10 | 0.00 | 0.01 | 2.14 |
| Factor 11 | 0.01 | 1.29 | 0.01 |
| Factor 12 | 0.02 | 1.14 | 0.02 |
| Factor 13 | 0.03 | 0.43 | 0.44 |
| Factor 14 | 0.01 | 0.78 | 0.10 |
| Factor 15 | 0.03 | 0.72 | 0.05 |
| Factor 16 | 0.03 | 0.66 | 0.06 |
| Factor 17 | 0.03 | 0.49 | 0.23 |
| Factor 18 | 0.02 | 0.56 | 0.03 |
| Factor 19 | 0.02 | 0.43 | 0.02 |
| Factor 20 | 0.01 | 0.38 | 0.02 |

**Supplementary Table 3. Genes associated to the top 20 CpGs in Factor 16.**

| **CpG ID** | **Gene ID** | **Location** | **Link** |
| --- | --- | --- | --- |
| cg19283806 | CCDC102B | (+) chr18:66382446-66722426 | https://ngdc.cncb.ac.cn/ewas/browse?probeId=cg19283806 |
| cg16867657 | RP1-62D2.3 | (+) chr6:11043757-11078459 | https://ngdc.cncb.ac.cn/ewas/browse?probeId=cg16867657 |
|  | ELOVL2 | (-) chr6:10980992-11044547 |  |
| cg19235307 | MBD4 | (-) chr3:129149787-129159022 | https://ngdc.cncb.ac.cn/ewas/browse?probeId=cg19235307 |
|  | IFT122 | (+) chr3:129158879-129239350 |  |
| cg02046143 | IGSF9B | (-) chr11:133778459-133826880 | https://ngdc.cncb.ac.cn/ewas/browse?probeId=cg02046143 |
| cg07082267 | GSE1 | (+) chr16:85203131-85709810 | https://ngdc.cncb.ac.cn/ewas/browse?probeId=cg07082267 |
| cg09027133 | MAN1C1 | (+) chr1:25943959-26112698 | https://ngdc.cncb.ac.cn/ewas/browse?probeId=cg09027133 |
| cg10501210 | AL035209.2 | (-) chr1:207974863-208052441 | https://ngdc.cncb.ac.cn/ewas/browse?probeId=cg10501210 |
| cg06240854 | TSGA10 | (-) chr2:99613724-99771427 | https://ngdc.cncb.ac.cn/ewas/browse?probeId=cg06240854 |
|  | RP11-111H13.1 | (+) chr2:99757948-99939204 |  |
|  | LIPT1 | (+) chr2:99771418-99779620 |  |
|  | RP11-38C17.1 | (+) chr2:99771461-99811761 |  |
| cg09124496 | AC005027.4 | (+) chr7:41733514-41818986 | https://ngdc.cncb.ac.cn/ewas/browse?probeId=cg09124496 |
|  | INHBA | (-) chr7:41706766-41745432 |  |
| cg18303397 | MBD4 | (-) chr3:129149787-129159022 | https://ngdc.cncb.ac.cn/ewas/browse?probeId=cg18303397 |
|  | IFT122 | (+) chr3:129158879-129239350 |  |
| cg23078123 | RP11-518D3.1 | (+) chr1:68297986-68668670 | https://ngdc.cncb.ac.cn/ewas/browse?probeId=cg23078123 |
|  | WLS | (-) chr1:68564156-68698803 |  |
| cg18933331 | No matching records found | | |
| cg07080372 | CMB9-55F22.2 | (+) chr11:797511-799190 | https://ngdc.cncb.ac.cn/ewas/browse?probeId=cg07080372 |
|  | SLC25A22 | (-) chr11:790475-798333 |  |
| cg20426710 | ZBED4 | (+) chr22:50247490-50283726 | https://ngdc.cncb.ac.cn/ewas/browse?probeId=cg20426710 |
| cg16585234 | MAN1C1 | (+) chr1:25943959-26112698 | https://ngdc.cncb.ac.cn/ewas/browse?probeId=cg16585234 |
| cg16932827 | No matching records found | | |
| cg23186333 | CD44 | (+) chr11:35160417-35253949 | https://ngdc.cncb.ac.cn/ewas/browse?probeId=cg23186333 |
| cg21406967 | SLC12A9 | (+) chr7:100424442-100464631 | https://ngdc.cncb.ac.cn/ewas/browse?probeId=cg21406967 |
|  | TRIP6 | (+) chr7:100464760-100471076 |  |
| cg01252023 | CORO1B | (-) chr11:67202981-67211292 | https://ngdc.cncb.ac.cn/ewas/browse?probeId=cg01252023 |
| cg13385220 | LGR6 | (+) chr1:202163029-202288909 | https://ngdc.cncb.ac.cn/ewas/browse?probeId=cg13385220 |
